# Changes in the Active, Dead, and Dormant Microbial Community Structure Across a Pleistocene Permafrost Chronosequence

**DOI:** 10.1101/457259

**Authors:** Alex Burkert, Thomas A. Douglas, Mark P. Waldrop, Rachel Mackelprang

## Abstract

Permafrost hosts a community of microorganisms that survive and reproduce for millennia despite extreme environmental conditions such as water stress, subzero temperatures, high salinity, and low nutrient availability. Many studies focused on permafrost microbial community composition use DNA-based methods such as metagenomic and 16S rRNA gene sequencing. However, these methods do not distinguish between active, dead, and dormant cells. This is of particular concern in ancient permafrost where constant subzero temperatures preserve DNA from dead organisms and dormancy may be a common survival strategy. To circumvent this we applied: (i) live/dead differential staining coupled with microscopy, (ii) endospore enrichment, and (iii) selective depletion of DNA from dead cells to permafrost microbial communities across a Pleistocene permafrost chronosequence (19K, 27K, and 33K). Cell counts and analysis of 16S rRNA gene amplicons from live, dead, and dormant cells revealed how communities differ between these pools and how they change over geologic time. We found clear evidence that cells capable of forming endospores are not necessarily dormant and that the propensity to form endospores differed among taxa. Specifically, Bacilli are more likely to form endospores in response to long-term stressors associated with permafrost environmental conditions than members of Clostridia, which are more likely to persist as vegetative cells over geologic timescales. We also found that exogenous DNA preserved within permafrost does not bias DNA sequencing results since its removal did not significantly alter the microbial community composition. These results extend the findings of a previous study that showed permafrost age and ice content largely control microbial community diversity and cell abundances.

**Importance:** The study of permafrost transcends the study of climate change and exobiology. Permafrost soils store more than half earth’s soil carbon despite covering ∽15% of the land area (Tarnocai et al 2009). This permafrost carbon is rapidly degraded following thaw (Tarnocai C et al 2009, Schuur et al 2015). Understanding microbial communities in permafrost will contribute to the knowledge base necessary to understand the rates and forms of permafrost C and N cycling post thaw. Permafrost is also an analog for frozen extraterrestrial environments and evidence of viable organisms in ancient permafrost is of interest to those searching for potential life on distant worlds. If we can identify strategies microbial communities utilize to survive permafrost we can focus efforts searching for evidence of life on cryogenic cosmic bodies. Our work is significant because it contributes to an understanding of how microbial life adapts and survives in the extreme environmental conditions in permafrost terrains across geologic timescales.

## Introduction

Permafrost contains active microbial communities that are moderately diverse (1) (2). DNA-based methods, such as metagenomics and 16S rRNA gene sequencing, are commonly used to interrogate these communities with the underlying assumptions that the data represent intact viable cells or that nonviable cells do not strongly affect conclusions drawn from whole community DNA. However, DNA from dead cells may drastically alter estimates of diversity and abundance. Furthermore, because many communities host dormant cells, DNA-based approaches may not represent active members. In temperate soils, up to 40% of DNA is from dead or compromised cells (3). In permafrost, the amount of ‘relic’ DNA may be even higher because frozen conditions preserve DNA from dead cells. In non-permafrost environments, multi-omic approaches provide functional information from RNA and protein, largely overcoming the problems of dormancy and relic DNA. In permafrost, this strategy has only been successfully applied in young near-surface permafrost due to low biomass and activity in older samples (4) (5). Therefore, in older deeper permafrost, other methods must be used to differentiate between live, dead, and dormant cells.

Alternative methods to multi-omics investigations, include microscopy, stable isotope probing, and physiological measurements with microbial isolates, have been used on permafrost samples to demonstrate that an active community exists. Electron-microscope examinations have shown evidence of apparently intact (no visual damage to cell envelopes), compromised (cell envelope ruptures), and dormant (endospores and cells with thick capsules) cells in permafrost (6) (7) (8). Using live/dead differential staining coupled with fluorescence microscopy, Hansen et al (2007) estimated that 26% of cells from a permafrost microbial community on Svalbard were viable (2). Stable isotope probing revealed that permafrost microorganisms can build biomass and replicate their genomes at subzero temperatures as demonstrated through the incorporation of ^14^ labeled acetate into lipids (9) and DNA (10). Similarly, studies involving permafrost microbial isolates have discovered microorganisms capable of reproduction at −15°C and metabolism down to −25°C (11) (12).

Though these studies show permafrost microbes exist in active states, dormancy is still a viable strategy for many taxa. Microorganisms enter dormancy in variety of ways, though the hardiest and most persistent is the endospore formed by some Gram-positive taxa in response to nutrient limitation, temperature extremes, or other stressors (13). However, endospore formation does not appear to be a universal survival strategy in permafrost because the relative abundance of endospore-forming taxa varies substantially across the Arctic and sub-Arctic, ranging from vanishingly rare to almost 80% (14) (15) (16). The abundance may be related to soil physicochemical properties including depth (17) (18), ice content, and permafrost age (16) (19). Furthermore, endospore-forming taxa in permafrost are not necessarily dormant. Hultman et al (2015) used RNA to DNA ratios to show that Firmicutes in young Holocene permafrost are more active than expected based on DNA abundance alone, demonstrating that dormancy cannot be inferred based solely on 16S rRNA gene amplicon sequencing (4).

While endospore formation may contribute to long-term survivorship in permafrost it is unclear whether this strategy is optimal across geologic timescales. Despite resistance to extreme conditions, DNA within an endospore can still accumulate damage (20). Typically, DNA damage is repaired upon germination by DNA repair machinery (13). However, damage accumulated over geologic timescales may be beyond the ability of repair enzymes to remedy. Willerslev et al (2004) amplified 16S rRNA genes from globally distributed permafrost soils ranging from 0 - 600 kyr old and found non-endospore-forming Actinobacteria were more highly represented than the endospore-forming Firmicutes in samples of increasing age (>100 kyr) (19). Johnson et al (2007) used a uracil-N-glycosylase treatment to break down damaged DNA extracted from ancient permafrost and found Actinobacteria rather than endospore-forming Firmicutes were more highly represented in the oldest samples (>600 kyr) (21). This suggests metabolic activity and active DNA repair may be a better survival strategy than dormancy in increasingly ancient permafrost. Therefore, endospore formation may not be an optimal survival strategy in permafrost for timescales beyond the Late Pleistocene.

Here we address the unresolved question of whether indicators of life (DNA, spores) in increasingly ancient permafrost are from live viable microbial communities or, instead, originate from only dead and dormant cells. We hypothesized that dormancy would increase with age but endospore formation wouldn’t be the sole mechanism for survival. Viable non-endospore forming taxa should also be present and some endospore-forming cells should exist in a non-dormant state. We also hypothesized that cell abundance would decrease with age, but cell abundance could also be affected by the biophysical conditions of the site when it was frozen into permafrost. Further, we asked whether preserved DNA from dead cells is so compositionally dissimilar from DNA from live cells that it alters inferences about microbial community structure in permafrost made with total soil DNA.

## Materials & Methods

### Permafrost Sample Collection

We collected frozen permafrost samples from the United States Cold Regions Research and Engineering Laboratory (CRREL) Permafrost Tunnel research facility located 16 km north of Fairbanks, Alaska (64.951°N, - 147.621°W) (Figure S1A). The facility includes 300 m of tunnels excavated into the permafrost. The tunnel where samples were collected extends 110 m horizontally at a depth of ∽15 m into a hillside exposing a chronosequence of late Pleistocene permafrost (Figure S1B) (22) (23). The temperature of the tunnel is maintained by refrigeration at −3 °C. It contains massive ice wedges (24) (25) surrounded by high organic content ice cemented windblown silt (26). In April 2016, we collected ice cemented silt from three locations inside the tunnel representing three age categories: 19K (approximately 10 m from the portal), 27K (54 m), and 33K (88 m) as determined previously by radiocarbon dating (16). After removing the sublimated surface layer (∽5 cm) from the walls of the tunnel, we collected five replicate cores per age category using a 7.5 × 5 cm key hole saw attached to a power drill as described previously (16). Cores were shipped back to California State University, Northridge (CSUN) on dry ice and stored at −20 °C.

### Permafrost Subsampling

For subsampling, we placed cores on autoclaved foil at room temperature for 10 minutes to allow the outer ∽1 cm to thaw and soften. Surface contamination was removed by scraping the outer layer with an autoclaved knife to expose the uncontaminated frozen interior. We sub-sectioned the remaining uncontaminated material using a fresh knife into sterile 50 ml falcon tubes and high-density polyethylene bags in preparation for downstream treatment.

### Soil Chemistry

Ice content was measured via gravimetric moisture analysis. To determine pH, soil was diluted 1:1 in a CaCl_2_ solution and measured using a Hannah benchtop meter with attached probe (Hanna Instruments, Woonsocket, RI). Percent total carbon, organic carbon, total nitrogen, and carbon:nitrogen were measured via dry combustion and direct measurement of total nutrients using an Elementar analyzer (Elementar, Langenselbold, Germany). Dissolved organic carbon (DOC) was measured using diluted meltwater on a Shimadzu total organic carbon (TOC) analyzer (Shimadzu Corporation, Kyoto, Japan). Electrical conductivity (EC) was measured using a digital benchtop meter with a potentiometric probe submerged in a diluted soil solution (Hanna Instruments, Woonsocket, RI).

### Cell Separation for Enumeration via Microscopy

For cell enumeration, cells were separated from the permafrost soil matrix using Nycodenz density cushion centrifugation as described previously (27) (28) (29) (30) (31). To separate cells from soil debris we disrupted 1.5 g of soil in a mild detergent consisting of 2 ml of 0.05% Tween 80 and 50 mM tetrasodium pyrophosphate buffer (TTSP) (28) (32) and sonicated for 1 minute at 20V using a QSonica ultrasonicator (QSonica, Newtown, CT) with a 0.3 cm probe. Sonicated samples were centrifuged at 750 x g for 7 min at 4 °C to remove large particles and debris. We extracted 600 μl of the supernatant and layered it over 600 μl of 1.3 g/L Nycodenz (Accurate Chemical, Westbury, NY) solution in a 2 ml tube. The tubes were centrifuged at 14,000 x g for 30 min at 4 °C. We transferred 600 μl of the upper and middle phase containing bacterial cells into a sterile 2 ml tube and centrifuged at 10,000 x g for 15 min at 4 °C. The supernatant was discarded, and the pellet was resuspended in 1 ml of 0.85% NaCl solution.

### Live/Dead Staining

The Live/Dead BacLight Bacterial Viability Kit (Invitrogen Detection Technologies, Carlsbad, CA) was used to differentially stain live and dead cells. We added 3 μl of a 1:1 mixture of 3.34 mM SYTO 9 and 20 mM propidium iodide solution to 1 ml cell suspensions as per the manufacturer’s protocols. Stained suspensions were incubated at room temperature for 15 minutes in the dark to allow the dyes to permeate cells and bind to DNA.

### Dapi Staining

DAPI staining was performed to obtain total cell counts. After removal of soil debris (as described above in Cell Separation for Enumeration via Microscopy), we added 3 μl of 14.3 mM DAPI stock solution to each 1 ml cell suspension. Stained suspensions were incubated in the dark at room temperature for 15 minutes.

### Cell Enumeration

We diluted and vacuum filtered the stained suspensions onto a 25 mm diameter 0.2 μm pore size black polycarbonate membrane which we placed on a slide with sterile forceps. Samples were observed at 100X magnification on a single focal plane using a Zeiss Axio Imager M2 fluorescence microscope coupled to an Apotome 2.0 System with appropriate filters for each stain (Zeiss, Oberkochen, Germany). We counted fifteen fields of view for live/dead and DAPI stained cells for each sample (33) (34). The average number of cells per field of view was multiplied by the area of the filter and the dilution factor then corrected for dry weight to calculate the average number of cells per gram of dry weight.

### Cell Separation for DNA Extraction (SHMP Method)

While Nycodenz density centrifugation is effective at removing soil debris, making it ideal for microscopic visualization, it is biased against endospores and heavily attached cells (35). To address this we used a second less-biased method to extract cells for DNA-based analyses (36). We disrupted 5 g of sample in 25 ml of 1% sodium hexametaphosphate buffer (SHMP) and sonicated for 1 minute at 20 V using a QSonica ultrasonicator with a 0.6 cm probe. The samples were left for 15 minutes to allow large particles and debris to settle before transferring the supernatant to a clean 50 ml tube. We added 15 ml of 1% SHMP to the pellet and sonicated again with a 0.6 cm probe for 1 minute at 20V. The mixture was incubated for another 15 minutes to allow debris to settle and then we combined the new supernatant with the supernatant from the previous step. To further remove large particles and debris we centrifuged the combined supernatant at 20 x g for 2 minutes. We then divided the supernatant equally into two 50 ml tubes as an experimental group (which received either the propidium monoazide treatment or the lysozyme enzyme treatment) and a control group (which received no treatment). These tubes were centrifuged at 10,000 x g for 15 minutes to pellet biomass. The biomass pellet was either stored at −20 °C to await DNA extraction (in the case of the control pellets) or immediately used for downstream treatments.

### Depletion of DNA from Dead Cells via Propidium Monoazide Treatment

To deplete DNA from dead cells we treated cells extracted using the SHMP method with propidium monoazide (PMAxx, Biotium Inc., Hayward, CA), which is a DNA-intercalating dye similar to the nucleic acid dye propidium iodide (37). It is selectively permeable, passing through the impaired membranes of dead cells, but it is unable to penetrate the membranes of living cells. In the presence of intense bright light, the azide group enables propidium monoazide to covalently cross link double stranded DNA, preventing its amplification via PCR (38).

For the propidium monoazide treatment we resuspended the extracted cell pellets in 500 μl of 0.85% NaCl solution and placed them in clear 1.5 ml microcentrifuge tubes. We added 2.5 μl of 20 mM propidium monoazide solution to each microcentrifuge tube resulting in a final concentration of 100 μM. We increased the concentration from the commonly used 50 μM due to the presence of leftover soil debris following cell extraction as recommended for environmental samples by Heise et al (2016) (39) (40). Tubes were incubated in the dark at room temperature for 10 minutes. After incubation we placed the tubes on a sheet of foil in an ice bucket to prevent warming. A 500 W halogen work lamp was placed 20 cm above the samples for 15 minutes. Every five minutes, we mixed the samples gently to ensure even light distribution. Following light exposure, we centrifuged samples at 10,000 x g for 15 minutes and discarded the supernatant. We stored these propidium monoazide treated pellets at −20 °C until use in downstream DNA extractions.

Cell extraction from soil fails to remove all soil particles, which can subsequently block light penetration and prevent propidium monoazide from cross-linking to DNA. To verify that our propidium monoazide treatment is effective in the presence of the small number of remaining particles we extracted cells from temperate control soils collected from the California State University, Northridge (CSUN) campus, spiked the sample with 3.6 x 10^8^ isopropanol-killed *Escherichia coli*cells, and treated the mixture with propidium monoazide. We also performed the treatment without *E. coli* spike-ins on the same temperate control samples. The amount of DNA removed was determined by comparing the number of copies of the 16S rRNA gene in the spiked and non-spiked samples before and after treatment (as determined by qPCR—see below for detailed protocols). The number of 16S rRNA gene copies decreased by ∽66% after treatment (Student’s t-test, t (8.64) = 3.6, p < 0.01) showing that even in the presence of soil particles, treatment removed 66% of added exogenous DNA.

### Endospore Enrichment via Lysozyme Enzyme Treatment

To separate endospores from vegetative cells we used a lysozyme enzyme treatment involving three steps: physical, enzymatic, and chemical cell lysis (hereafter referred to as lysozyme enzyme treatment following the convention of Wunderlin et al. (2016)) (36). The first physical treatment uses heat to lyse vegetative cells. Second, lysozyme dissolves the cell membrane followed by a solution of sodium hydroxide (NaOH) and sodium dodecyl sulfate (SDS) to further disrupt cellular membranes. Finally, a DNAse treatment is used to degrade the DNA from ruptured cells.

We resuspended cell pellets extracted using the SHMP method with 900 μl of 1X Tris - EDTA buffer (10 mM Tris and 1 mM EDTA; pH 8) and placed them into 2 ml tubes. Tubes were placed in a heat block at 60 °C for 10 min with shaking at 80 rpm. After incubation, we let the tubes cool for 15 min to 37 °C before adding 100 of lysozyme solution (20 mg/ml in 1X TE Buffer) and incubating in a heat block at 37 °C for 60 min with shaking at 80 rpm. After lysis 250 μl of 3N NaOH and 100 μl of 10 % SDS was added to the sample, reaching a final volume of 1.35 ml, which we incubated at room temperature for 60 min at 80 rpm. After the final incubation, we centrifuged the solution at 10,000 x g for 15 minutes to pellet cell debris and discarded the supernatant. We resuspended the pellet with 450 μl of water, 50 μl of 1X DNAse reaction buffer and 1 μl DNAse enzyme (New England Biolabs, Ipswich, MA) and left it for 15 min to remove DNA from lysed and dead cells. Following the DNase treatment we centrifuged the tubes for 15 min at 10,000 x g to pellet endospores and discarded the supernatant. The pellet was then resuspended in 1 ml 0.85% NaCl solution to wash away residual DNAse enzyme. We centrifuged the suspension at 10,000 x g for 15 min, discarded the supernatant, and stored the lysozyme enzyme treated pellet at −20°C in a sterile Eppendorf tube until it was used for downstream DNA extraction.

To confirm that lysozyme treatment depletes vegetative cells we extracted cells from temperate control soils collected on the CSUN campus, spiked the samples with 1.4 × 10^7^ live *E. coli* cells, and performed the lysozyme enzyme treatment. We used the same treatment on the temperate control soils without *E. coli* spike-ins. By comparing the number of copies of 16S rRNA genes before and after treatment in the spiked and non-spiked samples we were able to measure the amount of DNA removed (as determined by qPCR—see below for detailed protocols). Endospore enrichment treatment resulted in significant (82%) removal of DNA from vegetative *E.coli* cells (Student’s t-test, t (8.02) = −9.68, p < 0.01), demonstrating the effectiveness of lysozyme enzyme treatment in removing vegetative cells.

### DNA Extraction

We performed DNA extractions using a modified bead-beating protocol capable of lysing endospores, cysts, and cells with thickened walls, all of which are known to exist in permafrost (8) (42) (43). We resuspended propidium monoazide treated, lysozyme enzyme treated, and control pellets in 775 µl of lysis buffer (0.75M sucrose, 20 mM EDTA, 40 mM NaCl, 50 mM Tris) and transferred them to a MP Bio Lysis Matrix E Tube. We added 100 μl of 20 mg/ml lysozyme and incubated the samples at 37°C for 30 minutes. Following incubation, 100 µl 10% SDS was added and the samples were homogenized in an MP Biomedicals FastPrep 24 homogenizer for 20 seconds at 4.0 m/s. We placed the samples in a heat block at 99 °C for 2 minutes and allowed them to cool at room temperature for 5 minutes. We added 25 of 20 mg/ml Proteinase K and incubated samples at 55 °C overnight. The next day we centrifuged the tubes at 10,000 x g for 15 minutes and transferred the supernatant to a clean 2 ml Eppendorf tube. We used a FastDNA Spin Kit for Soil (MP Biomedicals, Santa Ana, CA) for DNA extractions from the lysed cells following the manufacturer’s protocols but omitting the initial lysis step. The DNA was cleaned using a PowerClean DNA Clean-Up (Mo Bio, Carlsbad, CA) and quantified using a Qubit 2.0 fluorometer (Thermofisher Scientific, Canoga Park, CA).

### PCR Amplification and Sequencing of the 16S rRNA gene

The variable region four (V4) of bacterial and archaeal 16S rRNA genes from the propidium monoazide treated, lysozyme enzyme treated, and control groups was amplified as described by Caporaso et al (2012) but with the addition of 2 µl of 20 mg/ml bovine serum albumin in each PCR reaction. We used the golay barcoded primer set 515F/806R with an added degeneracy to enhance amplification of archaeal sequences on the 515F primer and thermal cycling steps recommended by the Earth Microbiome Project protocol version 4.13 (44). The amplified PCR products from triplicate reactions for each sample were pooled at approximately equal concentrations, as measured using a PicoGreen dsDNA Quantification Assay Kit (Thermofisher Scientific, Canoga Park, CA). We quantified the pooled 16S rRNA gene amplicons by qPCR using an Illumina Library Quantification Kit (Kapa Biosystems, Wilmington, MA) on a CFX96 Real Time PCR Detection System (Bio-Rad, Hercules, CA). 16S rRNA gene amplicons were sequenced with a 2 × 150 bp v2 Reagent Kit on an Illumina MiSeq instrument.

We demultiplexed and quality filtered raw fastq data using the Quantitative Insights Into Microbial Ecology (QIIME) software package version 1.9.1 (45).Sequences were truncated at the first position with a quality score less than 3. Then forward and reverse sequences were merged with a minimum merged sequence length of 200 bp, a minimum overlapping length of 20 bp, and a maximum of one difference in the aligned sequence. All sequences that passed quality filtering were *de novo* clustered into operational taxonomic units (OTUs) at 97% sequence identity using USEARCH (46).We assigned taxonomy using the RDP classifier (47) with a confidence score of 0.5 (48) (49). For phylogenetic metrics of diversity, a phylogenetic tree was constructed using FastTree (50) as implemented in QIIME. We rarefied samples to an equal depth (N =5000 sequences/sample) for all subsequent analyses. One OTU was only abundant (<3% relative abundance) in the blank samples and samples which had the lowest DNA yields. This OTU, from the genus Burkholderia, was removed from all other samples because it was likely a result of laboratory contamination (51). 16S rRNA gene sequence data were uploaded to the NCBI’s sequence read archive under accession number SRP158034.

### Quantitative PCR of the 16S rRNA gene

qPCR of the V4 region of the 16S rRNA gene was conducted on propidium monoazide treated and lysozyme treated temperate soil DNA after receiving a spike in of *E. coli* to evaluate these methods. Amplification was accomplished using the 515F/806R primer set. Triplicate qPCR reactions were done in 25 μl volumes (12.5 μl of GoTaq qPCR Master Mix (Promega, Madison, WI), 1.25 μl of each primer (17) (44), 5 μl nuclease free water, and 5 μl of template DNA) in 96 well plates on a CFX96 Real Time PCR Detection System (Bio-Rad, Hercules, CA). The thermal cycler program was as follows: 95 °C for 2 min, 40 cycles of (95 °C 15 sec, 60 °C 60 sec), and a melt curve analysis (60-95 °C). Quantified full-length *E. coli* 16S rRNA gene amplicons were used to make a standard curve. A negative control lacking template DNA was performed along with each qPCR.

### Statistical Analysis

We tested for differences in soil chemical characteristics between age categories using a Kruskal Wallis and a Dunn’s post hoc test for nonparametric data using the ‘PMCMR’ package in R (52). P-values were corrected using the False Discovery Rate (FDR). Significant differences in the ratio of live to dead cells, the proportion of live cells, and the direct cell counts for each stain between age categories was also tested using the same methods.

We used the Shannon Diversity Index and phylogenetic diversity metrics to calculate alpha diversity using the ‘alpha_diversity.py’ script in QIIME (53). Differences in alpha diversity between age categories and treatments were tested the same methods as for soil chemistry. We computed beta diversity using the weighted UniFrac metric (54). Differences between samples were visualized using principal coordinate analysis (PCoA) using the ‘phyloseq’ package in R (55). We also used non-metric multidimensional scaling coordinates of microbial community composition between samples for PerMANOVA analysis as a function of soil chemistry data using the function ‘adonis’ from the ‘vegan’ package in R (54) (56).

Differences in the relative abundance of specific taxa between treatments and age categories were tested using a linear mixed effect model on rank transformed taxa abundances and nested treatment as a factor within age using the ‘nlme’ package in R (57). P-values were corrected using the FDR. The specific taxa indicated were tested for various pairwise comparisons between treatments and untreated controls for each age category using a Mann-Whitney-Wilcoxon test using the ‘wilcoxon.test’ function in R. Differences in 16S rRNA gene copy number, as quantified using qPCR, between propidium monoazide treated samples and untreated controls (including samples given a spike in with dead *E. coli* cells) and the endospore enriched samples (given a spike in with live *E. coli* cells) were tested with a t-test using the ‘t.test’ function in R.

## Results

### Soil Chemistry

Soil physicochemical properties including ice content, carbon, nitrogen, pH, and conductivity varied significantly among age categories (Kruskal Wallis, p > 0.01 for all measurements, Table 1). These values were consistently higher in the intermediate age category compared to the older and younger samples.

### Cell Enumeration

We performed cell counts across the chronosequence using live/dead and DAPI staining coupled with fluorescent microscopy. Average cell counts ranged from 3.6 × 10^6^ to 9.2 × 10^6^ cells [gram dry weight (gdw)^-1^] (live cells), 1.7 × 10^7^ to 4.5 × 10^7^ cells gdw^-1^ (dead cells), and 2.3 × 10^7^ to 4.7 × 10^7^ cells gdw^-1^ (total cell count). Live (Kruskal Wallis, X^2^ (2) = 46.25, p < 0.001), dead (Kruskal Wallis, X^2^ (2) = 53.16, p < 0.001), and total (Kruskal Wallis, X^2^ (2) = 53.58, p < 0.001) counts were significantly different among categories and higher in the intermediate age samples compared with the oldest (Dunn’s test, p < 0.01) and youngest (Dunn’s test, p < 0.01) samples (Figure 2A). The proportion of live cells was significantly higher in intermediate aged samples (26%) compared to the youngest samples (14%) (Kruskal Wallis, X^2^ (2) = 9.84, p < 0.01, Dunn’s test, p < 0.01, Figure 2B). In the oldest samples, 25% of the cells were live though this was not significantly different than the values observed for the youngest or intermediate aged samples. The ratio of live cells to dead cells did not change significantly across the chronosequence (Kruskal Wallis, X^2^ (2) = 1.38, p > 0.05).

**Figure 1.**
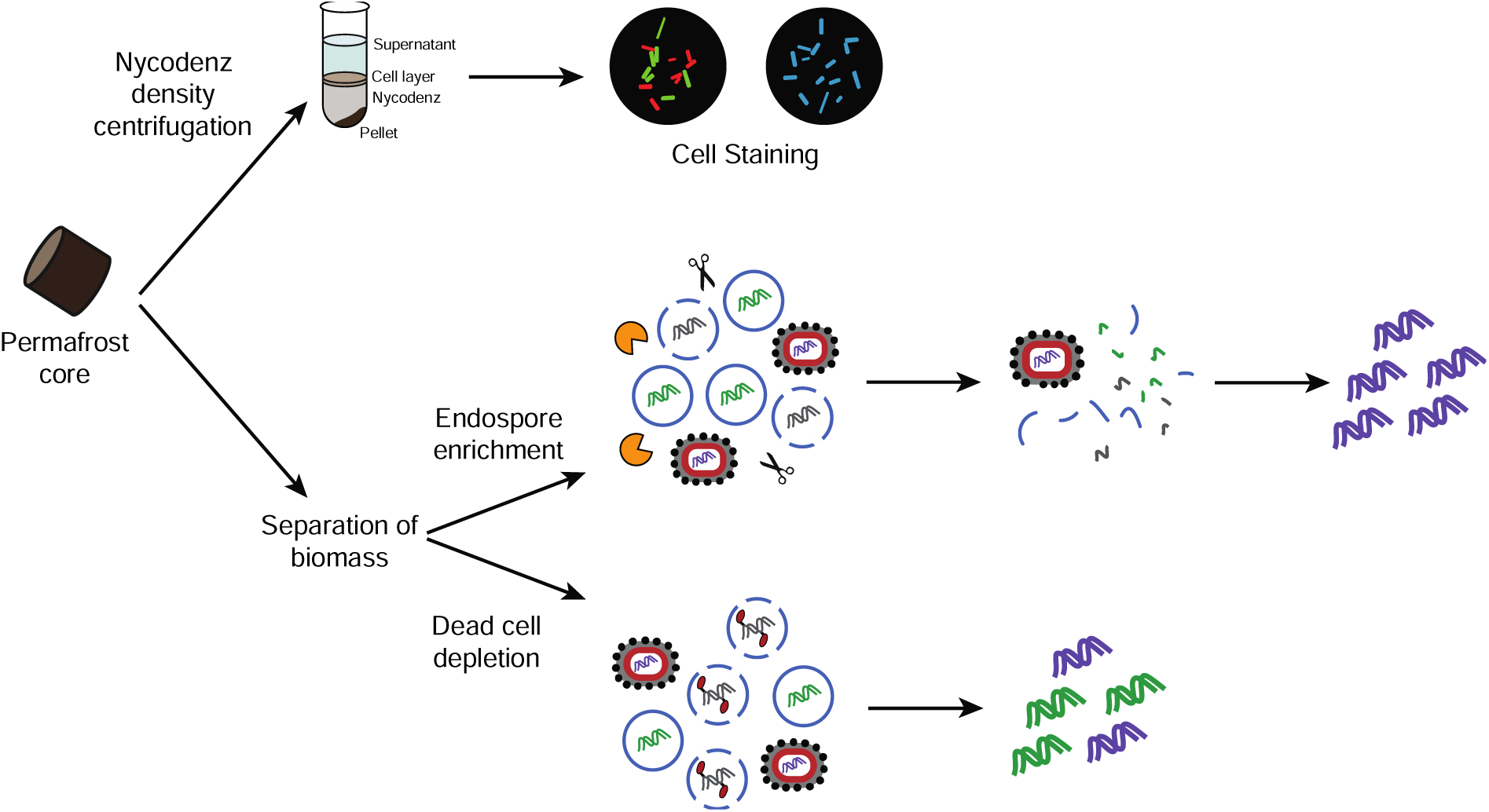
Experimental strategy overview. Live, dead, and dormant cell counts were conducted by separating cells from soil using a Nycodenz density centrifugation, staining with a Live/Dead differential stain or DAPI, and counting via fluorescence microscopy. For the endospore enrichment and dead cell depletion experiments, we separated biomass from soil using a gravity separation technique. To deplete DNA from dead organisms, cell mixtures were treated with propidium monoazide and then exposed to light, causing cross-links with DNA not enclosed by an intact cell envelope or spore coat. The crosslinks inhibit downstream PCR amplification. To enrich for endospores, cell mixtures were exposed to lysozyme, heat, and DNAse, which lyses vegetative cells and degrades DNA. In both the endospore enrichment and dead cell depletion experiments, the 16S rRNA gene was amplified and used for downstream analysis.

**Figure 2.**
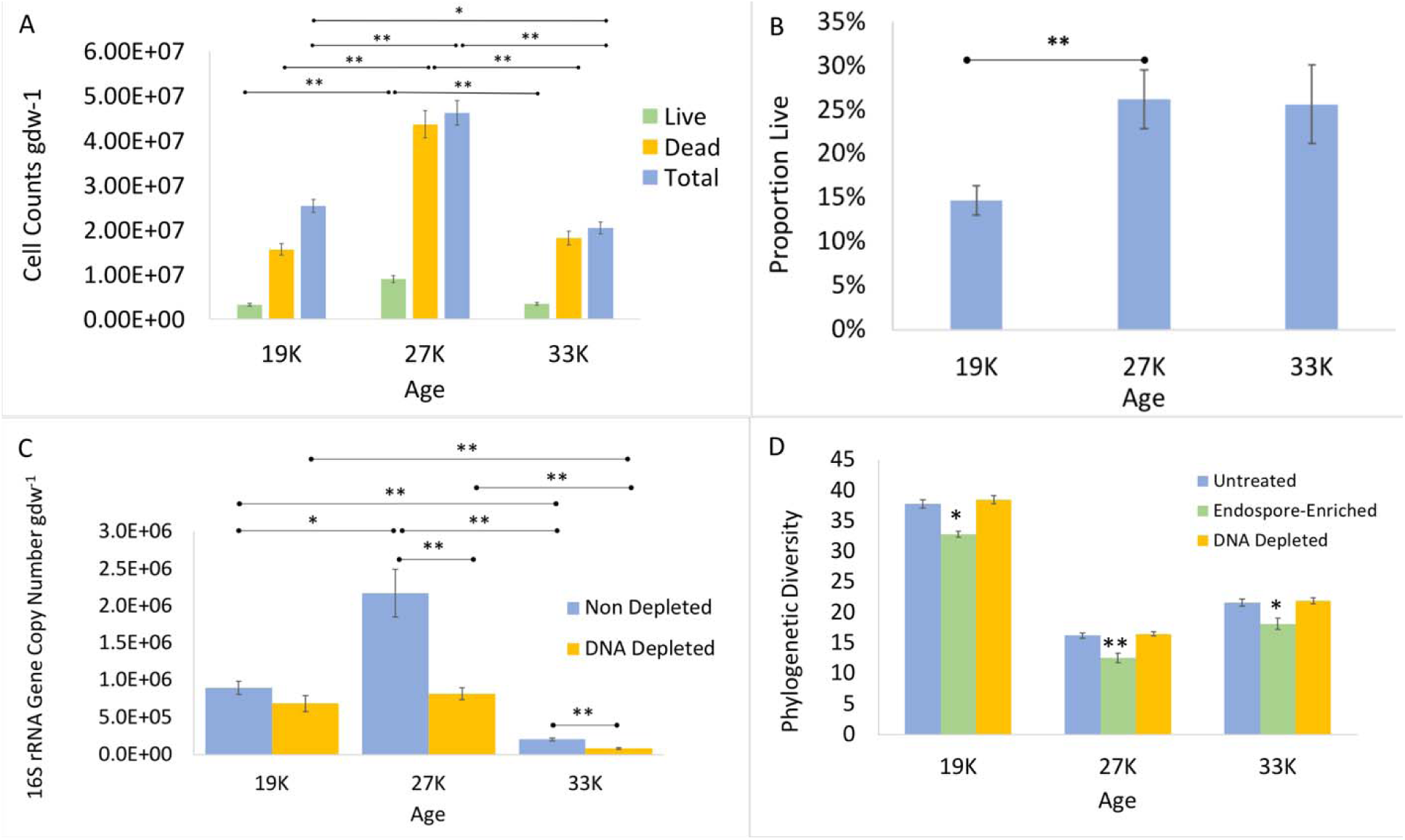
(A) Direct cell counts as determined by cell staining and fluorescent microscopy. Live/dead and total (DAPI) counts. (B) Proportion of live cells as determined by direct counts of live cells (stained with SYTO 9) and total cells (stained with DAPI). (C) 16S rRNA gene copy number in samples depleted of dead DNA using propidium monoazide and non-depleted controls. qPCR of the V4 region of the 16S rRNA gene in samples showed that average copy number decreased for all age groups in the propidium monoazide treated group (DNA depleted samples) compared to the untreated group (non-depleted controls) and that there was a decrease in viable DNA across the three age categories. (D) Phylogenetic diversity index compared across age categories and treatment types. Values show the average of five replicate cores and error bars show the standard error of the mean. Significant p-values tested by Dunn’s post-hoc test are indicated (* p < 0.05, ** p > 0.01).

### Endospore Enrichment via Lysozyme Enzyme Treatment

The community of cells remaining after lysozyme enzyme treatment had reduced alpha-diversity compared with non-treated controls as measured via phylogenetic richness (Kruskal Wallis, p < 0.05,Figure 2D) and the Shannon Index (Kruskal Wallis, p < 0.05, Figure S2). The enzyme treatment increased the relative abundance of three phyla—Firmicutes, Actinobacteria, and Chlamydiae (Figure 3A). Endospore enrichment tended to increase the relative abundance of Firmicutes in all age categories but was only significant for the youngest (Mann-Whitney-Wilcoxon test, U = 0, p < 0.01) and intermediate (Mann-Whitney-Wilcoxon test, U = 2, p < 0.05) age categories. At the class level, endospore enrichment changed the abundance of Bacilli and Clostridia, but in opposing directions. It increased the relative abundance of Bacilli across all age categories, growing more pronounced in the older samples (youngest: 5.8%, intermediate: 9.3%, oldest: 18.6%). These data were significant for each age category (Mann-Whitney-Wilcoxon test, U = 2, p < 0.05, Figure 3B). In contrast, endospore enrichment significantly decreased the relative abundance of Clostridia in the oldest age category (10%) (Mann-Whitney-Wilcoxon test, U = 2, p < 0.05, Figure 3B). Treatment did not significantly change the relative abundance of Clostridia in the youngest and intermediate samples. These trends were driven by the families *Planococcaceae, Thermoactinomycetaceae, Bacillaceae,* and *Paenibacillaceae* for Bacilli and the family *Clostridiaceae* for Clostridia (Table 2).

**Figure 3.**
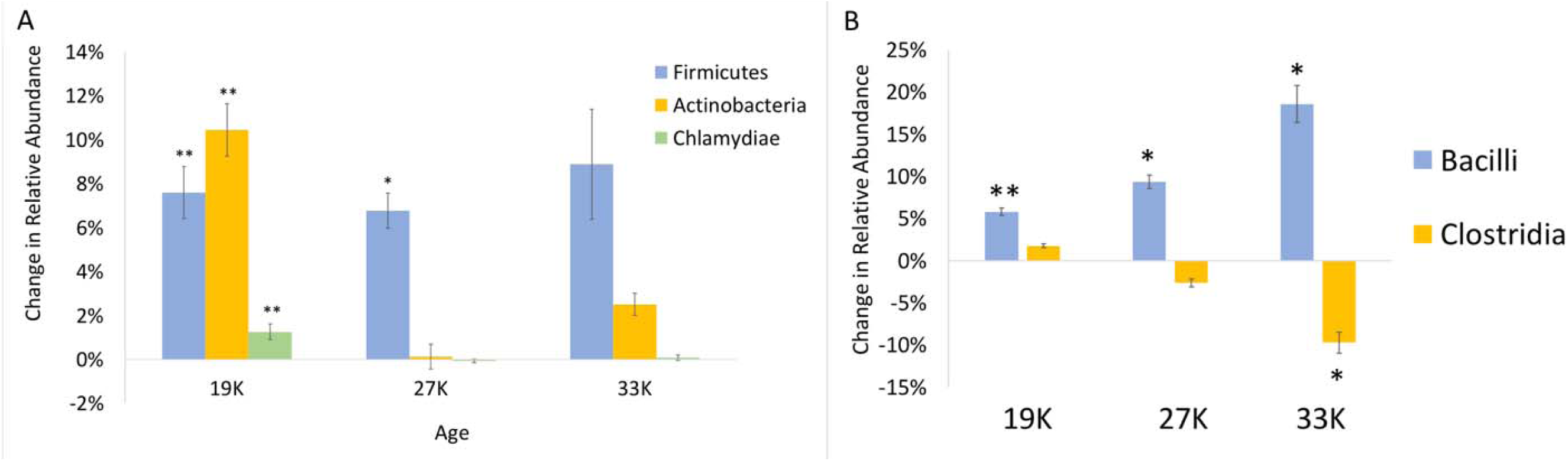
(A) Phylum-level changes in relative abundance due to endospore enrichments. (B) Class-level changes in relative abundance due to endospore enrichments. Values show averages of five replicate coress and error bars show standard error of the mean. Significant p-values tested by the Mann-Whitney-Wilcoxon test are indicated (* p < 0.05, ** p > 0.01).

**Table 1.**
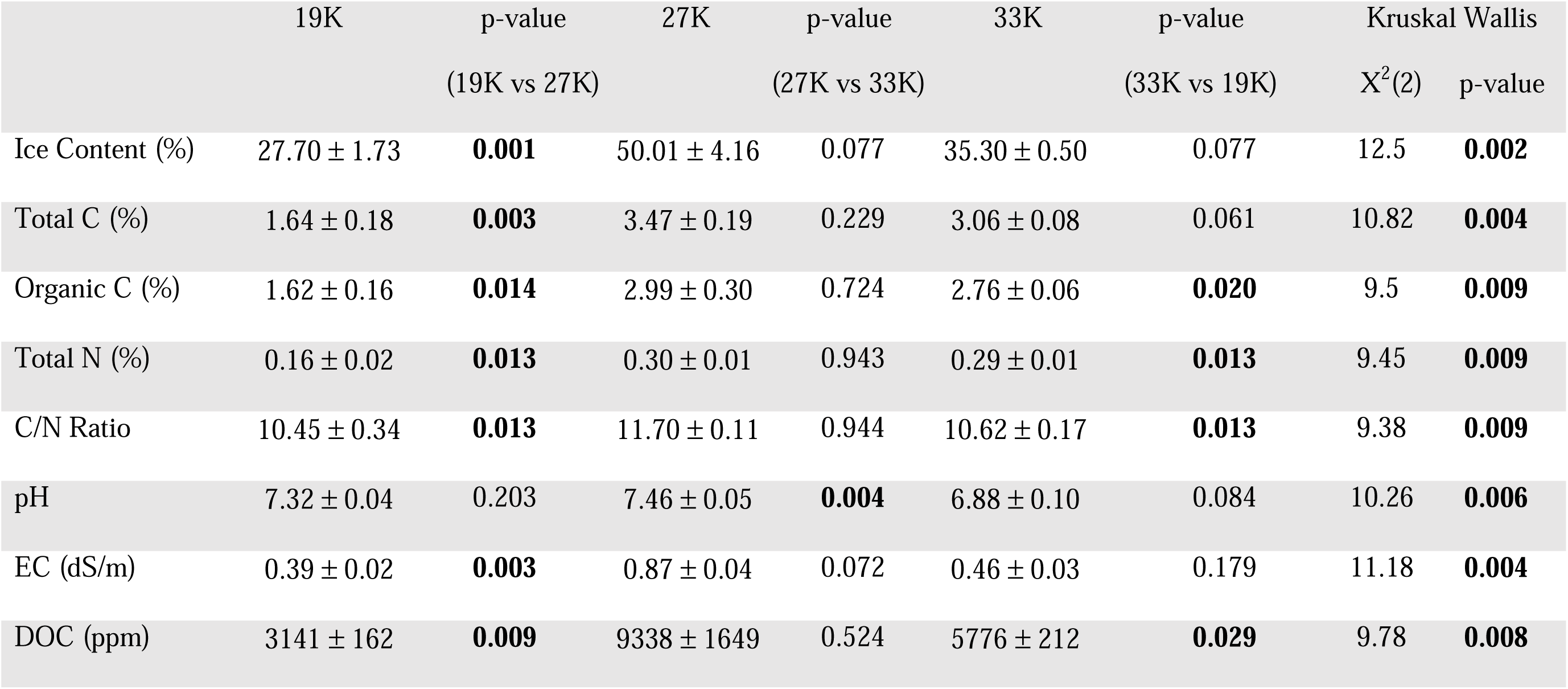
Permafrost physicochemical characteristics across the three time periods. Values are averages of five replicates plus/minus one standard error of the mean. Statistical differences were tested using a Kruskal Wallis test and a Dunn’s post hoc test with p-values corrected using the False Discovery Rate. Significant p-values (p < 0.05) are show in bold.

**Table 2.** Average percent difference in relative abundance between the endospore-enriched samples and non-enriched controls across the three age categories. A negative value shows underrepresentation in the endospore-enriched samples compared to the nonenriched controls while a positive value shows overrepresentation. Values show averages of five replicate cores.

| Taxa | Family | 19K (%) | U-value | 27K (%) | U-value | 33K (%) | U-value |
| --- | --- | --- | --- | --- | --- | --- | --- |
| Actinobacteria | *Micrococcaceae* | 2.4 ± 0.5 ** | 0 | 0.0 ± 0.0 | 9.5 | 0.0 ± 0.0 | 11 |
|  | *Solirubrobacteraceae* | 1.5 ± 0.4 | 3 | 0.6 ± 0.3 | 4 | 0.1 ± 0.1 | 5 |
|  | *Gaiellaceae* | 1.0 ± 0.3 * | 1 | -0.2 ± 0.1 | 8 | 0.1 ± 0.1 | 8.5 |
| Bacilli | *Planococcaceae* | 2.4 ± 0.9 | 4 | -8.5 ± 1.0 * | 1 | 4.0 ± 2.6 | 8 |
|  | *Thermoactinomycetaceae* | 0.4 ± 0.1* | 0 | 20 ± 2.5 * | 2 | 7.8 ± 2.2 | 6 |
|  | *Bacillaceae* | 0.7 ± 0.1 * | 0 | -2.5 ± 1.2 | 8 | 1.0 ± 0.4 | 9 |
|  | *Paenibacillaceae* | 2.5 ± 0.4 * | 0 | 0.4 ± 1.6 | 12 | 5.6 ± 2.7 | 4 |
| Clostridia | *Clostridiaceae* | 3.1 ± 0.4 ** | 0 | -2.5 ± 1.5 | 5 | -9.2 ± 1.2 ** | 0 |

(Mann-Whitney-Wilcoxon test, \*\* $p < 0.01$ , \* $p < 0.05$ ).

Endospore enrichment increased the relative abundance of Actinobacteria in the youngest age category from 22.4% to 32.8% (Mann-Whitney-Wilcoxon test, U = 0, p < 0.01, Figure 3A). This increase was driven by the families *Micrococcaceae* within the Actinomycetales, as well as the *Gaiellaceae* and *Solirubrobacteraceae* (Table 2). There were no significant differences in Actinobacteria relative abundance due to endospore enrichment in the intermediate and oldest samples.

Chlamydiae relative abundance increased in the youngest age category from 0.7% to 2.0% as a result of endospore enrichment (Mann-Whitney-Wilcoxon test, U = 0, p < 0. 01, Figure 3A). This trend was driven by an increase in the relative abundance of the family *Chlamydiaceae.* All other major taxa including Proteobacteria, Alphaproteobacteria, Deltaproteobacteria, Bacteroidetes, Acidobacteria, Chloroflexi, and Planctomycetes decreased in relative abundance due to the endospore enrichment (Table S1).

### Depletion of DNA from Dead Cells via Propidium Monoazide Treatment

To determine if DNA from dead cells biases estimates of taxonomic relative abundance from the whole community we used a propidium monoazide treatment to deplete dead cell DNA. Depletion did not significantly change the relative abundance of taxa at the phylum, class, order, or family levels for any of the age categories (Mann-Whitney-Wilcoxon test, p > 0.05). Actinobacteria were consistently less abundant in depleted samples compared with non-depleted controls. Unlike the results from endospore-enrichment experiments, propidium monoazide treatment did not change alpha diversity measurements—there were no significant differences in depleted samples compared with non-depleted controls (Kruskal Wallis, p > 0.05, Figure 2D & S2).

To confirm propidium monoazide treatment successfully depleted DNA from dead and membrane-compromised cells we determined 16S rRNA gene copy number in treated and untreated samples using qPCR. Depletion decreased copy number for the youngest (∽24%, Student’s t-test, p > 0.05), intermediate (∽62%, Student’s t-test, t (16) = 4.11, p <0.01), and the oldest samples (∽58%, Student’s t-test, t (26) = 5.45, p < 0.01, Figure 2C). 16S rRNA gene copy number varied significant across ages for the depleted samples (Kruskal Wallis, X^2^ (2) = 25.05, p < 0.001) and control samples (Kruskal Wallis, X^2^ (2) = 31.25, p < 0.001). The oldest samples had significantly fewer 16S rRNA gene copies compared to the youngest (Dunn’s test, p < 0.01) and intermediate (Dunn’s test, p < 0.01) for both the depleted and control samples (Figure 2C).

### 16S rRNA gene-based community analysis

To explore how experimental treatments influenced overall community structure we compared 16S rRNA gene sequences from 19K, 27K, and 33K old permafrost from treated and control samples (n = 5 replicate samples x 3 age categories x 4 treatments = 60 samples total). Comparisons of diversity between samples revealed that samples clustered by age regardless of treatment (Figure S3A). When comparing the samples within each age category, endospore enriched samples consistently clustered separately from controls and DNA-depleted samples (Figure S3B-D). Phyla that differed significantly based on age and treatment (endospore enrichment and depletion of DNA from dead cells) using a linear mixed effect model included Proteobacteria, Actinobacteria, Firmicutes, Bacteroidetes, Chloroflexi, Acidobacteria, Planctomycetes, and Chlamydiae (FDR corrected p < 0.01).

Overall, alpha diversity estimates were highest in the youngest category and lowest in the intermediate age (Table S2). All soil chemistry measurements correlated significantly with non-metric multidimensional scaling vectors (PerMANOVA, p < 0.05, Figure S4). Age and ice content were the strongest drivers of community structure (PerMANOVA, R^2^ = 0.49, F = 181.35, p < 0.001; PerMANOVA, R^2^ = 0.26, F = 91.98, p < 0.001). Other measurements had a small but significant effect on community structure: DOC (PerMANOVA, R^2^ = 0.04, F = 13.13, p < 0.001), C:N (PerMANOVA, R^2^ = 0.03, F = 12.27, p < 0.001), Total C (PerMANOVA, R^2^ = 0.02, F = 6.63, p < 0.01), and Total N (PerMANOVA, R^2^ = 0.01, F = 3.43, p < 0.05).

## Discussion

In this study we used three complimentary methods—two of which have never been applied to permafrost—to distinguish between live, dead, and dormant cell types: live/dead staining, endospore enrichment, and depletion of DNA from dead cells. We present evidence that endospore-forming organisms observed in DNA-based studies are not always dormant. Instead, among potential endospore-forming organisms the tendency to exist as an endospore was taxa dependent. Under the stressful conditions associated with long-term interment in permafrost, members of class Bacilli were more likely to exist as endospores in increasingly ancient permafrost while members of Clostridia were more likely to remain in a non-dormant state. This trend grew more pronounced as sample age increased. For Bacilli, this trend was driven by the families *Planococcaceae*, *Thermoactinomycetaceae, Bacillaceae,* and *Paenibacillaceae* which are predominantly aerobes (58). For Clostridia, this trend was driven by the family *Clostridiaceae* which decreased by over 9% in the endospore-enriched samples compared to non-enriched controls from the oldest aged category. Most members of the family *Clostridiaceae* are obligate anaerobes, with metabolic strategies ranging from fermentation to anaerobic respiration to chemolithoautotrophy (58). The limited oxygen availability in permafrost could be a stressor that contributes to endospore formation for groups of Bacilli while anaerobic and metabolically diverse Clostridia are able to persist in a non-dormant state.These data suggest endospore formation is a viable survival strategy for Bacilli against the conditions of ancient permafrost, at least for the timescales observed in this study. However, over increasing timescales endospores can accumulate DNA damage. In the absence of active repair machinery, this damage may become lethal (13) (19) (21). Further investigations into even older permafrost may show decreases in endospore forming taxa in favor of organisms that can actively repair DNA damage as has been suggested previously (19).

The presence of endospore-forming Firmicutes is highly variable across Arctic permafrost though they have been shown to increase in relative abundance over geologic timescales (16). Tuorto et al (2014) used stable isotope probing to identify active community members and found that Firmicutes were not among those replicating their genomes at subzero temperatures with the caveat that stable isotope probing only identifies taxa that actively replicate their genomes. The study would not have identified cells that were metabolically active but not dividing during the experimental timeframe. (10). Other studies demonstrate that endospore-forming taxa are active. For example, Hultman et al (2015) found the abundance of transcripts from Firmicutes exceeded expected levels based on the abundance of DNA sequence reads (4).

Though our endospore treatment was designed to enrich for true endospores it also enriched for two other phyla—Actinobacteria and Chlamydiae—but only in the youngest age category. Actinobacteria were previously found to resist this treatment perhaps due to their ability to form dormant and spore-like structures (59). In our samples, members of families *Micrococcaceae, Solirubrobacteraceae,* and *Gaiellaceae* were overrepresented in our endospore-enriched group compared to non-enriched controls. Members of these families can survive radiation, starvation, and extreme desiccation (21) (58), which may also confer the ability to resist lysozyme treatment.

Unexpectedly, Chlamydiae were also more abundant in the endospore-enriched samples compared to non-enriched controls from the youngest age category. This was driven exclusively by the family *Chlamydiaceae,* which are not known to have resting states (58). All genera within this class are obligate intracellular symbionts of Acanthamoeba. Acanthamoeba are a genus of single-celled eukaryotes commonly found in freshwater and soil. They exist in free-living forms and as stress-resistant dormant cysts (60) which could account for the increase in the relative abundance of Chlamydiae in the endospore-enriched samples. Several studies have found intact and viable Acanthamoeba cysts in Holocene and Pleistocene permafrost (61) (62). Though we did not sequence eukaryotic marker genes, it is possible that there are Acanthamoeba in our samples.

Depleting DNA from dead cells did not significantly alter microbial community composition suggesting that sequencing 16S rRNA marker genes from total DNA provides a reasonable representation of community structure in permafrost. This result would be expected when death rate and the rate of degradation of dead cells is proportional among taxa (63). Though depletion experiments did not bias estimates of relative abundance, it reduced 16S rRNA gene copy number by ∽48%, suggesting that ∽1/2 of DNA from permafrost is ‘relic DNA.’ Our estimate of relic DNA is higher than values taken from temperate soil where ∽40% of DNA is relic (3).

One potential limitation of using the propidium monoazide treatment to deplete DNA from dead cells is that soil particles can prevent light penetration and limit efficacy.We attempted to mitigate this concern by extracting biomass from soil particles, performing intermittent mixing during treatment, and increasing PMA concentration for particle-rich samples (39). We also confirmed the efficacy of the experiment by spiking samples with dead *E. coli* cells and showed that even in the presence of soil particles, treatment removed 66% of added exogenous DNA.

Direct cell counts and analysis of 16S rRNA amplicons from permafrost soils showed that age and ice content were the most important drivers of cell abundance and microbial diversity, similar to a prior study conducted in the Permafrost Tunnel with samples collected directly adjacent ours (16). While endospore enrichment significantly altered microbial community structure within each age group compared to non-enriched samples, the effects of age and ice content were still much stronger predictors of beta diversity among samples from all ages.

Microbial cell counts ranged from 2.3 10^7^ to 4.7  10^7^ cells gdw^-1^ and were consistent with previous studies from Arctic and sub-Arctic permafrost (1) (14) (15) (64) (65), but were one to two orders of magnitude lower than is commonly observed in temperate soils (66) (67) (68). Among the three age categories the intermediate had the highest number of live, dead, and total cells and the highest proportion of live cells. 16S rRNA gene copy number from total community DNA and from samples depleted of dead DNA using propidium monoazide showed the highest counts in the intermediate age category and the lowest counts in the oldest category, similar to cell count data derived from live/dead staining. The oldest samples also had a markedly higher proportion of live cells than the youngest samples, though the data narrowly missed the significance cutoff. The high proportion of live cells in the intermediate and oldest samples may be attributable to the higher levels of DOC measured in those samples. It also may be due to increased reliance on necromass as labile carbon becomes less available in older samples. This is supported by metagenomic evidence from Permafrost Tunnel microbes, showing an increase in the abundance of genes related to scavenging detrital biomass in older permafrost (16).

Though cell counts were highest in the intermediate age category, alpha diversity was lower compared with the youngest and oldest samples. The decrease in alpha diversity may reflect the high ice content in those cores, which has been shown previously to decrease diversity in permafrost (42) (69). The observation that cell counts are greatest in the intermediate samples is consistent with a previous publication from the Permafrost Tunnel chronosequence, though counts here are greater by an order of magnitude. This is likely due to increased cell recovery as a result of our biomass extraction protocol, which used sonication (rather than vortexing) to separate cells from soil particles (16).

The Live/Dead staining approach uses membrane permeability as a proxy for viability (70) and has been used extensively in environmental samples (71) (72) (73) including permafrost (2) (16). One potential drawback to using this approach is that live cells can incorrectly stain as dead, particularly under dark anoxic conditions (74). We suggest this is unlikely to impact our samples because permafrost is a generally stable environment in which membrane potentials are well maintained (75) and we stained under aerobic conditions in the light. However, if our treatment affected membrane potentials, this would result in an underestimate of the number of live cells.

This study builds on previous work aimed at understanding how microbial communities adapt to the extreme conditions of permafrost over geologic timescales (16). We used two strategies never before applied to permafrost (endospore-enrichment and depletion of DNA from dead organisms) to show that Firmicutes represented in DNA based studies are not always dormant. We found that preserved DNA from dead organisms does not bias DNA sequencing results, since its removal using propidium monoazide did not significantly alter microbial community composition. We confirmed that both microbial cell counts and diversity decreased between our youngest and oldest samples and were primarily controlled by sample age and ice content. We also found that despite the multiple treatments used, differences in community composition among age categories were robust. There are changes within age categories caused by treatments but it suggests future studies interrogating microbial communities among diverse permafrost types may similarly show strong patterns driven by soil physiochemical properties, including age. Age-driven survival strategies and community structure identified in this and a companion study from the Permafrost Tunnel (16) may be common across other types of permafrost.

Expanding investigations to older permafrost samples and permafrost representing different biogeochemical properties will be essential to building a model for how microorganisms in permafrost survive for geologic timescales. The question of dormancy is a key building block of this model. While dormancy appears to be a survival strategy for microbes not well adapted to life in permafrost (e.g. Bacilli, which are commonly aerobes or facultative anaerobes), those that appear to be better adapted (e.g. Clostridia, which are typically anaerobes and have a diverse suite of metabolic strategies to draw upon) are less likely to resort to dormancy. This suggests an increase in endospore formation among maladapted taxa upon entrance to the late Pleistocene, but that metabolic activity may be increasingly necessary in older permafrost. Expanding the age range of permafrost and conducting further in-depth studies of current samples (such as quantitative measurements of endospore markers compared with vegetative cells markers), will be crucial. If life exists on cryogenic bodies in the solar system, it must have persisted for longer time scales than exist in Earth’s permafrost. Thus, a chronosequence-based approach to understanding survival may allow us to extrapolate to more ancient cryoenvironments.

Finally, permafrost communities may have an important role in the biogeochemical cycling of elements and greenhouse gas production following thaw. Our data suggest DNA-based studies provide a reasonable representation of the taxonomic reservoir present and poised to decompose soil carbon upon thaw. It also demonstrates that the taxonomic diversity and the number of cells is governed by soil physicochemical characteristics of permafrost soil. Communities with less diversity and fewer cells may be slower to respond to thaw, which has implications for the speed at which carbon is processed and released into the atmosphere. Our data suggests that the reservoir (both the number of cells and the diversity) of microbes is controlled by age, carbon, and moisture content and highlights the need for detailed understanding of how physicochemical properties shape microbial communities across permafrost environments.

## Acknowledgements

This work was supported by the National Aeronautics and Space Administration (NASA) (grant numbers NNX15AM12G and NNH15AB58I). AB acknowledges support from the California State University, Northridge Biology Department Graduate Student Tuition Waiver. We thank Steven Escalante, Tara Mahendrarajah David Romero, and Archana Srinivas for assistance with permafrost subsampling.

